# A chemical-genetic interaction between PAF1 and ENL/AF9 YEATS inhibition

**DOI:** 10.1101/2025.09.09.675200

**Authors:** Paige A. Barta, Leopold Garnar-Wortzel, Timothy R. Bishop, Rachel E. Hayward, Lauren M. Hargis, James B. Shaum, Hui Si Kwok, Brian B. Liau, Benjamin F. Cravatt, Michael A. Erb

**Affiliations:** Department of Chemistry, The Scripps Research Institute, La Jolla, CA, USA; Department of Chemistry and Chemical Biology, Harvard University; Cambridge, MA 02138, USA

## Abstract

Transcriptional regulatory proteins are frequent drivers of oncogenesis and common targets for drug discovery. The transcriptional co-activator, ENL, which is localized to chromatin through its acetyllysine-binding YEATS domain, is preferentially required for the survival and pathogenesis of acute leukemia. Small molecules that inhibit the ENL YEATS domain show anti-leukemia effects in preclinical models, which is thought to be caused by the downregulation of pro-leukemic ENL target genes. However, the transcriptional effects of ENL YEATS domain inhibitors have not been studied in models of intrinsic or acquired resistance and, therefore, the connection between proximal transcriptional effects and downstream anti-proliferative response is poorly understood. To address this, we identified models of intrinsic and acquired resistance and used them to study the effects of ENL YEATS domain inhibitors. We first discovered that ENL YEATS domain inhibition produces similar transcriptional responses in naive models of sensitive and resistant leukemia. We then performed a CRISPR/Cas9-based genetic modifier screen and identified in-frame deletions of the essential transcriptional regulator, PAF1, that confer resistance to ENL YEATS domain inhibitors. Using these drug-resistance alleles of *PAF1* to construct isogenic models, we again found that the downregulation of ENL target genes is shared in both sensitive and resistant leukemia. Altogether, these data support the conclusion that the suppression of ENL target genes is not sufficient to explain the anti-leukemia effects of ENL antagonists.

## Introduction

The transcriptional co-activator, ENL (eleven-nineteen leukemia, also known as MLLT1), has been identified as a critical acute leukemia dependency by multiple independent groups.^1,2^ ENL and its paralog, AF9, contain a conserved YEATS (Yaf9, ENL, AF9, Taf14, and Sas5) domain that binds to acetylated lysine side chains and mediates chromatin localization.^2–6^ Genetic experiments have previously pointed to the YEATS domain as a potential drug target for ENL-dependent leukemia,^1,2^ motivating multiple groups, including our own, to develop small-molecule chemical probes that competitively inhibit the binding of ENL/AF9 YEATS domains to acetyllysine side chains.^7–10^ In leukemia cells, ENL binds disproportionately to the promoter of leukemia proto-oncogenes, such as *HOXA9/10, MEIS1, MYC*, and *MYB*, and recruits higher-order transcriptional regulatory complexes to chromatin, including the super elongation complex (SEC), which contains the master regulator of transcription elongation, P-TEFb (a CDK9/CycT heterodimer).^1,2,11–16^ ENL/AF9 YEATS domain inhibitors selectively repress the transcription of leukemic ENL target genes by preventing the recruitment of the SEC to their promoters, which we and others have proposed as the underlying explanation for their anti-proliferative effects.^1,2,8,10^

The selective suppression of leukemia proto-oncogenes following ENL/AF9 YEATS domain inhibition is also observed following ENL degradation and *ENL* knockout, pointing toward a coherent set of on-target transcriptional effects related to ENL loss-of-function in leukemia.^1,2,17,18^ Likewise, the anti-proliferative effects are on-target for at least one ENL/AF9 YEATS domain inhibitor, TDI-11055, as validated by the discovery of mutant *ENL* alleles that confer resistance to its inhibition of leukemia growth.^10^ However, no prior studies have directly tested whether the transcriptional suppression of ENL target genes is necessary and sufficient for the growth effects of ENL antagonists. In studying small molecules that inhibit transcriptional regulatory proteins, we and others have, in many instances, failed to distinguish sensitive and insensitive cells by their transcriptional responses.^19–21^ For example, BET bromodomain inhibition causes identical primary transcriptional effects in models of leukemia that are sensitive and insensitive to the downstream anti-proliferative effects of these agents.^22^ We recently observed the same phenomenon for CBP/p300 acetyltransferase inhibitors, suggesting that the primary transcriptional effects of these compounds are not sufficient to explain their anti-proliferative effects.^21^ Here, we use models of naive and acquired resistance to study the relationship between the transcriptional effects of ENL/AF9 YEATS domain inhibitors and their downstream effects on leukemia growth and survival.

## Results and discussion

ENL antagonists are proposed to function by selectively suppressing the transcription of genes that are involved in promoting self-renewal and proliferation.^7,8,10^ While this has been observed repeatedly in preclinical models of acute leukemia that are sensitive to ENL loss-of-function,^8,10,18,23,24^ the transcriptional effects of ENL antagonists have not been characterized in models that are intrinsically resistant to these compounds. To systematically identify models of naively sensitive and resistant acute leukemia cells, we profiled the anti-proliferative effects of SR-0813, an ENL/AF9 YEATS domain inhibitor previously reported by our group,^8^ across ∼900 cell lines using the PRISM assay (profiling relative inhibition simultaneously in mixtures) (Fig. 1A,B and Table S1, ESI).^25,26^ We also analyzed the cancer dependency map (DepMap), a catalog of essential genes in thousands of cancer cell lines,^27^ finding in both datasets that AML and B-ALL cell lines are preferentially sensitive to loss of ENL function (Fig. 1B-D, Table S2, ESI). This is consistent with previous reports that acute leukemia driven by MLL-fusion oncoproteins are most sensitive to ENL loss-of-function, which we also observed by PRISM (Fig. S1A).^1,2^ We identified some false-negative results in these datasets, such as MOLM-13 cells, which previous studies have shown are sensitive to both *ENL* knockout and ENL/AF9 YEATS domain inhibition but did not register as being dependent on ENL in DepMap (Fig. 1D).^1,2,8,10^ In contrast, the PRISM assay identified MOLM-13 as being among the most sensitive cell lines in the dataset to SR-0813, consistent with prior studies using structurally distinct ENL/AF9 YEATS domain inhibitors, including SR-0813 and TDI-11055 (Fig. 1A,C).^8,10^ Altogether, the orthogonal PRISM and DepMap datasets were broadly convergent, providing an unbiased validation that SR-0813 preferentially targets ENL-dependent acute leukemia cell lines.

**Fig. 1.**
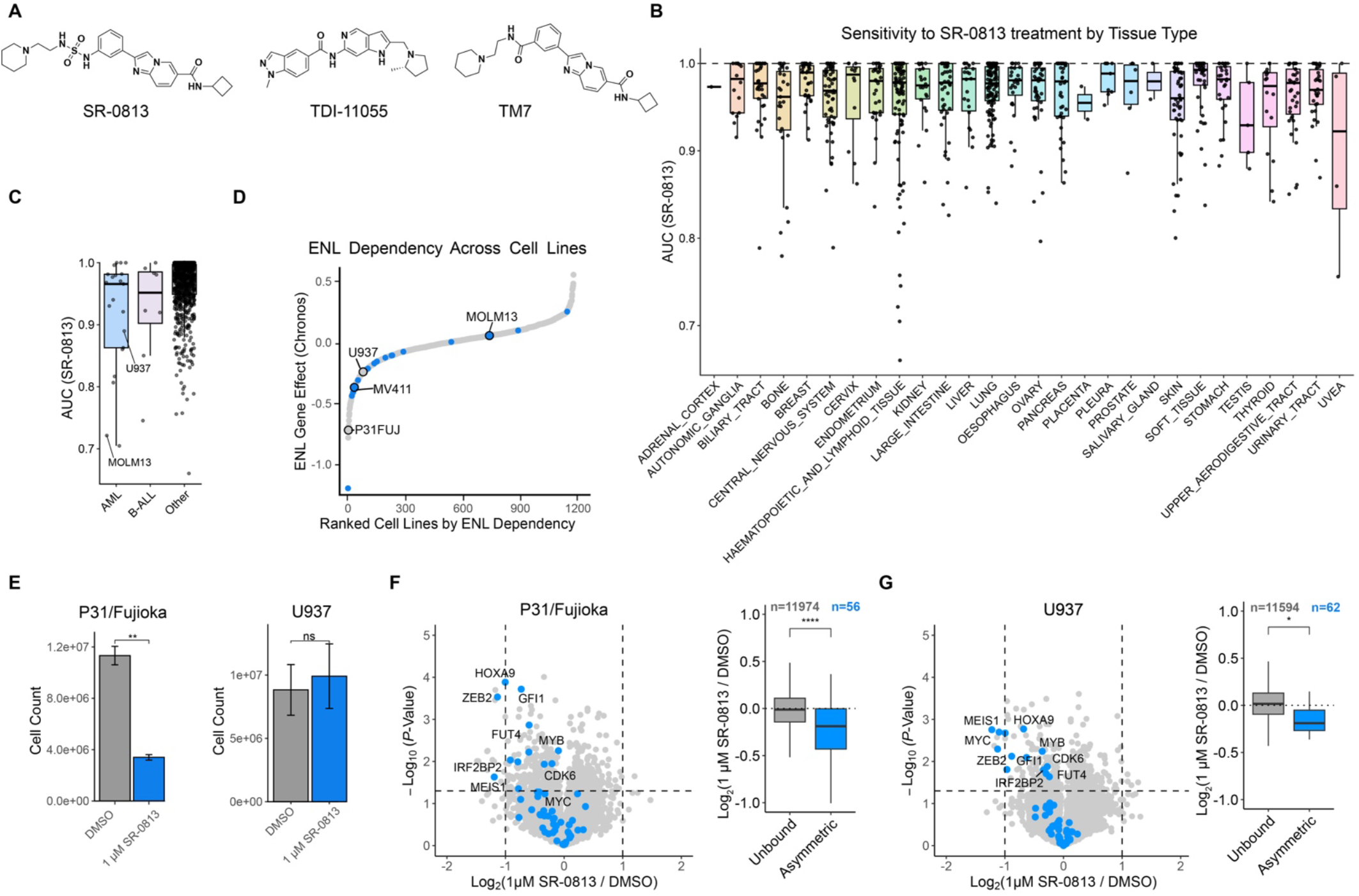
(A) ENL/AF9 Inhibitors used in this study. (B) PRISM (Profiling Relative Inhibition Simultaneously in Mixtures) data of SR-0813 sensitivity grouped by tissue. (C) PRISM data grouped by AML, B-ALL, and all other lineages. (D) Waterfall plot of ENL genetic dependency showing *ENL* gene effect scores across DepMap. Cell lines harboring MLL-rearrangements are shown in blue. (E) Cell growth counts after 11 days of treatment with DMSO or 1 µM SR-0813. (F) Volcano plot (left) and boxplot (right) of DMSO-normalized gene expression changes in P31/Fujioka cells in response to SR-0813 (4 h, 1 µM). ENL asymmetric target genes are shown in blue in the volcano plot and boxplot. Other genes that do not harbor ENL binding at their promoters (“unbound”) are shown in grey. Genes that are labeled are shared in both P31/Fujioka and U937 datasets. *P*-values are calculated with a Welch’s two sample *t*-test. (*P*-values: ^*^P<0.05, ^**^P<0.01, ^***^P<0.001, ^****^P<0.0001). (G) Same as in F for U937 cells.

Using the PRISM and DepMap datasets, we selected P31/Fujioka and U937 cells to study the transcriptional effects of ENL antagonism in sensitive and resistant leukemia models, respectively. Although both cell lines are driven by *CALM-AF10* fusions, they exhibit differential sensitivity to *ENL* knockout, providing lineage- and oncogene-matched models of sensitive and resistant acute leukemia (Fig. 1D). Of all cell lines in the DepMap dataset, P31/Fujioka cells are among the most sensitive to *ENL* knockout (Fig. 1D). In contrast, U937 are less sensitive to *ENL* knockout, and a previous study has demonstrated that they are insensitive to the ENL/AF9 YEATS domain inhibitor, TDI-11055.^10^ We confirmed these findings, showing that P31/Fujioka cells, but not U937 cells, are sensitive to SR-0813 (Fig. 1E).

To compare the underlying transcriptional effects of ENL/AF9 YEATS domain inhibition in these sensitive and resistant cell line models, we performed 3’-end mRNA-seq following 4 hours of SR-0813 treatment. In both models, SR-0813 significantly decreased the transcription of asymmetric ENL target genes, which are high-confidence ENL target genes marked by a disproportionate amount of ENL binding within the promoter region of a target gene (Fig. 1F,G and Tables S3,4, ESI).^1^ Notably, the proto-oncogenes *MYC, MYB, and HOXA9* were downregulated in U937, despite this cell line being insensitive to the anti-proliferative effects of SR-0813 (Fig. 1F,G). These data suggest that the preferential downregulation of ENL target genes may be insufficient to predict downstream growth responses.

To evaluate this possibility in isogenic models of sensitive and resistant leukemia, we sought to identify mechanisms of acquired resistance to SR-0813 using a genome-scale CRISPR/Cas9 screen in MOLM-13 cells. A gene-level analysis by MAGeCK (Model-based Analysis of Genome-wide CRISPR/Cas9 Knockout)^28^ revealed that sgRNAs targeting common essential genes (as defined by DepMap) were preferentially depleted in both the vehicle-treated (DMSO) and SR-0813-treated arms of the screen, confirming that the experiment performed as expected (Fig. 2A, Fig. S1B and Table S5, ESI). In querying for genes that were selectively enriched in the cells treated with SR-0813, we noted that the common essential gene *PAF1* (*RNA polymerase associated factor 1*) was one of the strongest hits (Fig. 2A).

**Fig. 2.**
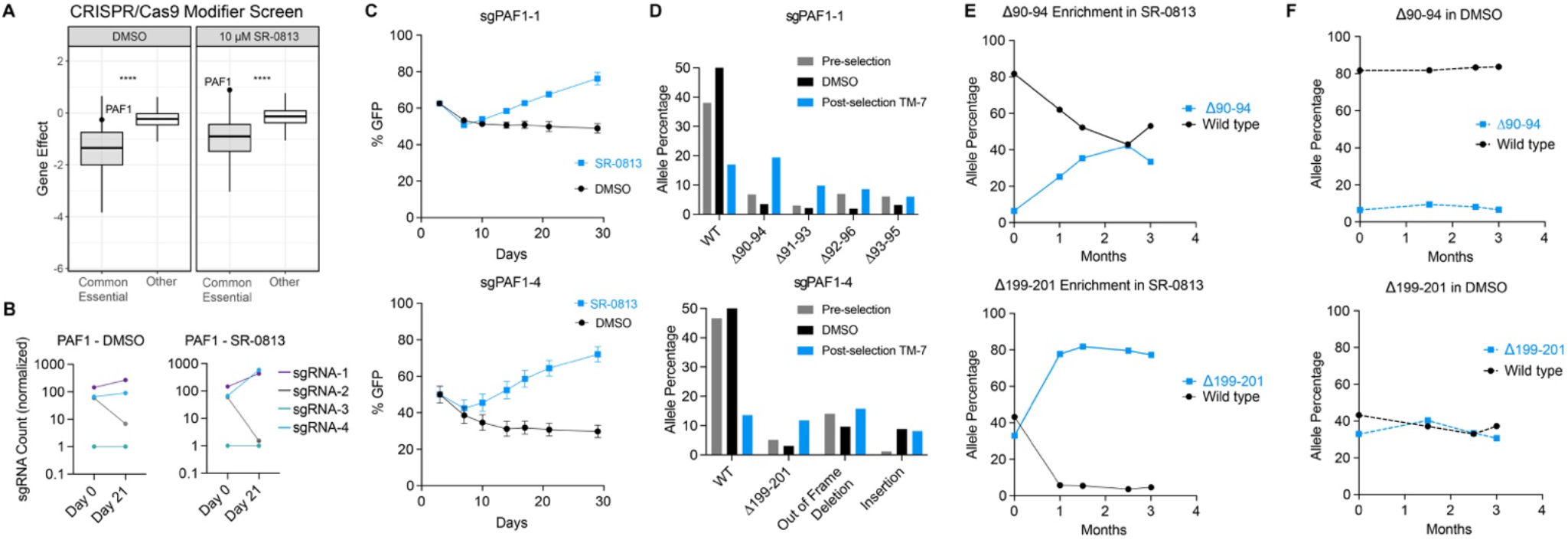
(A) MAGeCK MLE analysis of a pooled CRISPR/Cas9-based genetic screen in MOLM13-Cas9 cells comparing sgRNA proportion in DMSO- and SR-0813-treated cells (10 µM) after 21 day treatment time. (B) Enrichment of individual *PAF1* sgRNAs. (C) Competitive growth assay in MOLM13-Cas9 cells transduced with *PAF1*-targeting sgRNAs and treated with 10 µM SR-0813 or DMSO. (D) CRISPRESSO analysis of *PAF1* allele frequency following transductions with *PAF1* sgRNAs (pre-selection) and after growth in TM7 or DMSO for 30 days (post-selection). (E-F) *PAF1* alleles with in-frame deletions were introduced by CRISPR/Cas9-mediated homology-directed repair and tracked over time with nanopore sequencing during treatment with 10 µM SR-0813 (E) or DMSO (F).

Of the four *PAF1*-targeting sgRNAs included in the pooled CRISPR screen, two were enriched by SR-0813 treatment, which we termed sgPAF1-1 and sgPAF1-4 (Fig. 2B). We validated the effects of both sgRNAs in an arrayed format using competitive growth experiments. Here, MOLM-13 cells stably expressing Cas9 were transduced with a bicistronic vector encoding an sgRNA and GFP, such that the percentage of sgRNA-positive cells can be tracked over time by measuring the percentage of GFP-positive cells.^29^ MOLM-13 cells transduced with sgRNAs targeting *PAF1* were enriched in response to SR-0813 treatment but not by the DMSO control, validating the pooled screening results (Fig. 2C). We repeated this experiment with TDI-11055 and obtained similar results (Fig. S1C), indicating that *PAF1*-mediated resistance is a shared feature of ENL/AF9 YEATS domain inhibitors rather than a scaffold-specific effect of SR-0813.

PAF1 is a core component of the multiprotein PAF complex (PAFc), an evolutionarily conserved regulator of transcription by RNA Polymerase II (Pol II) with genetic and physical links to ENL/AF9 function.^30–37^ PAF1 has been shown to be required for leukemogenic transformation by *MLL*-fusion oncogenes and to interact directly with both MLL and ENL.^11,16,38–41^ Germline variants of *CTR9*, which encodes a subunit of PAFc, predispose to myeloid malignancies by losing the ability to antagonize the ENL/AF9-containing SEC in hematopoietic progenitors.^42^ Furthermore, both *ENL* and *CTR9* are recurrently mutated in Wilms tumors.^43–45^

Since *PAF1* is a common essential gene, we speculated that resistance to SR-0813 might be mediated by in-frame insertions or deletions that could preserve its role in cell survival.^27,46^ Previous studies have described in-frame indels resulting from genome-scale CRISPR screens, supporting this possibility.^47^ To identify the *PAF1* alleles that confer resistance to ENL/AF9 YEATS domain inhibition, we repeated the competitive growth experiment using a synthetically simplified analog of SR-0813, TM7,^17^ and collected the initial and final cell populations for nanopore-based sequencing of *PAF1*. We found that both sgRNAs created in-frame deletions of *PAF1* that were enriched in the population by TM7 treatment, leading to a concurrent loss of the wild-type *PAF1* allele (Fig. 2D). For sgPAF1-1, several partially overlapping deletions spanning amino acids 90-96 were enriched by TM7 treatment, with the strongest effect being observed for Δ90-94 (Fig. 2D). For sgPAF1-4, we observed a notable enrichment of a Δ199-201 allele.

To test whether these in-frame deletions are sufficient to confer resistance to ENL/AF9 YEATS domain inhibition, we introduced the Δ90-94 and Δ199-201 alleles into the endogenous *PAF1* locus in MOLM-13 cells using CRISPR/Cas9-based gene editing and homology-directed repair. Nanopore-based DNA sequencing of the edited populations over time showed that cells harboring these deletions were enriched by drug treatment while the wild-type sequence was depleted, indicating that these deletions confer resistance to ENL/AF9 YEATS domain inhibition (Fig. 2E,F). Notably, the PAF1 deletion affecting amino acid residues 199-201 exceeded an allele frequency of ∼80%, likely indicating that some percentage of cells survived in the absence of a wild-type allele.

PAF1 functions as a member of the PAF complex, which consists of 5 additional proteins: CTR9, LEO1, CDC73, WDR61 and RTF1. Using a previously reported cryo-electron microscopy (cryo-EM) structure (PDB 6TED), we found that the Δ90-94 and Δ199-201 deletions occur at sites of intracomplex protein-protein interactions, with Δ90-94 mapping to a site of contacts with CTR9 and Δ199-201 mapping to its interface with LEO1 (Fig. 3A,B, Fig.S2A,B).^48^ *CTR9* variants that predispose to myeloid malignancies were previously shown to cause partial loss of function by impairing PAF complex assembly.^42^Interestingly, the amino acid mutations identified in these *CTR9* variants occur along the interface of CTR9 and PAF1, with one of the identified mutations (N157S) in close proximity to PAF1 residues 90-94 (Fig. S2B,C).^42^Therefore, we speculated that in-frame PAF1 deletions might similarly impair the integrity of the PAFc and produce a state of hyperactive SEC activity that can compensate for the effects of ENL/AF9 YEATS domain inhibition.

**Fig. 3.**
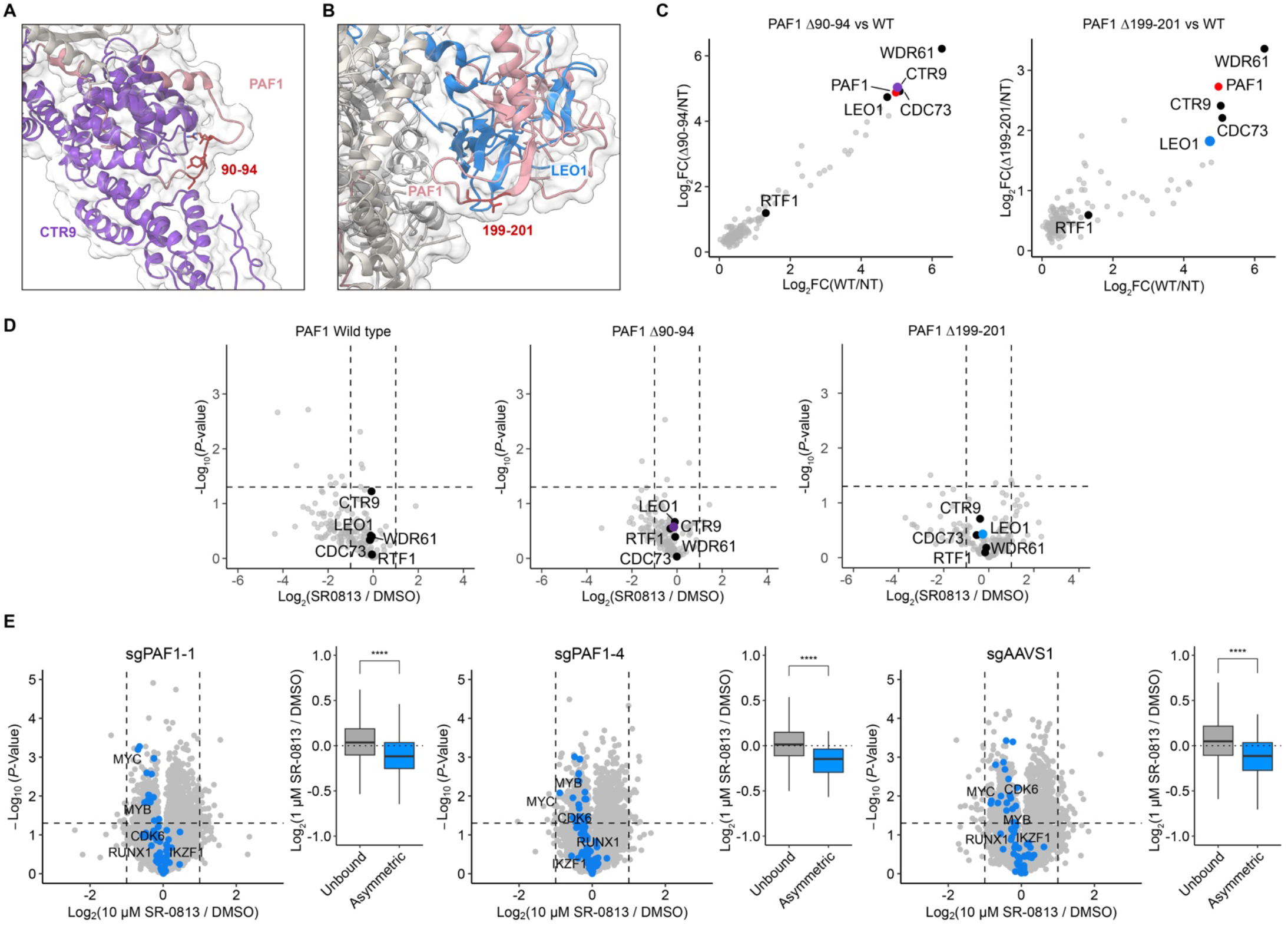
(A-B) Visualization of in-frame deletion sites within the PAF complex, as previously determined by cryo-EM (PDB 6TED). (C) Analysis of PAF1 IP-MS comparing wild-type (WT) and in-frame deletion alleles to the non-transduced (NT) control. (D) IP-MS comparing SR-0813 treatment with DMSO (normalized to PAF1 enrichment). (E) Volcano plots depicting gene expression changes in sgPAF1-1, sgPAF1-4, and sgAAVS1 cell lines upon treatment with SR-0813 (10 µM, 4 h) and boxplots comparing genes not bound by ENL (unbound, *n* = 12702) and asymmetric ENL target genes (*n* = 69). Labeled genes are those found across all three datasets. *P*-values are calculated with a Welch’s two sample *t*-test. (*P*-values: ^*^P<0.05, ^**^P<0.01, ^***^P<0.001, ^****^P<0.0001).

To address this, we performed co-immunoprecipitations followed by mass spectrometry (IP-MS) on MOLM-13 cells exogenously expressing epitope-tagged alleles of wild-type or mutant *PAF1*. This experiment showed clear enrichment of PAFc proteins but did not indicate that there was a significant loss of PAF1 interactions with corresponding complex members due to either deletion (Fig. 3C, Fig. S2C, Table S6, ESI). We also performed the co-IP experiments in the presence of SR-0813 but failed to observe any difference in the enrichment of PAF1c subunits in the presence of drug in wild-type or mutant cells (Fig. 3D). Although ENL is known to interact with PAF1, we did not detect ENL in our pulldown.

These data argue against our hypothesis that the Δ90-94 and Δ199-201 deletions in PAF1 affect PAFc interactions or stability. Nevertheless, we considered that they could still impact transcriptional responses to ENL YEATS domain inhibitors through an alternative mechanism. To assess this, we performed 3’-end mRNA sequencing on MOLM-13-Cas9 cells that were transduced with sgPAF1-1 or sgPAF1-4 and selected with SR-0813 over several weeks. Despite being resistant to the anti-proliferative effects of SR-0813, ENL target genes were similarly downregulated in cells expressing sgPAF1-1 and sgPAF1-4, compared to a control sgRNA targeting the *AAVS1* safe harbor locus (Fig. 3E, Table S7, ESI). We performed gene set enrichment analysis (GSEA), which further confirmed that ENL target genes are among the most substantially downregulated gene sets in both each dataset (Fig. S3A). The basal expression of ENL target genes prior to drug treatment was similar in each cell line, demonstrating that the downregulation of ENL targets is not compensated for by an overall increase in expression (Fig. S3B,C). Altogether, these data suggest that the direct downregulation of ENL target genes is likely insufficient to predict growth responses to ENL/AF9 YEATS domain inhibitors, consistent with our observations in U937 and P31/Fujioka cells.

## Concluding Remarks

Drug-resistant alleles provide exceptionally useful models for assessing the on- and off-target effects of small molecule drugs, offering an unbiased view of the biological pathways that control response and resistance.^49,50^ Here, we describe the discovery of mutant *PAF1* alleles with small, in-frame deletions that are sufficient to confer resistance to the antiproliferative effects of structurally diverse ENL/AF9 YEATS domain inhibitors. Surprisingly, we failed to detect a similar resistance to the transcriptional effects of ENL/AF9 YEATS domain inhibition, potentially suggesting that YEATS domain inhibitors may decrease proliferation in sensitive cancers *via* an alternative aspect of ENL/AF9 biology. Both ENL and PAF1 are reported to be involved in the regulation of DNA damage repair,^51–54^ pointing toward a possible area of ENL biology for future exploration with YEATS domain inhibitors.

Alternatively, it is possible that cellular adaptations, occurring secondary to an initial transcriptional effect, could control response and resistance to these compounds. For example, BET bromodomain inhibitors elicit similar primary transcriptional responses in sensitive and resistant cell lines, but the secondary adaptation to these changes have been shown to control response and resistance.^19,20,22^ We have observed parallels to this with CBP/p300 bromodomain inhibitors, where sensitive and resistant cell lines show similar primary transcriptional responses but starkly different transcriptional effects at later time points.^21^ Recently, global inhibitors of transcription, such as triptolide and Α−amanitin, were revealed to induce a form of programmed cell death, termed Pol II degradation-dependent apoptotic response (PDAR).^55^ In this context, cells rapidly activate apoptotic signaling in response to the loss of hypophosphorylated RNA Pol II, not as a result of changes in gene expression. Whether this mechanism could be involved in the response to ENL antagonists that suppress transcription in a gene-specific manner is unclear.

While the precise mechanism by which *PAF1* alterations confer resistance to YEATS domain inhibitors remains unresolved, our study provides strong support for the conclusion that the anti-proliferative effects of SR-0813 are on target. Notably, we found that *PAF1* deletion alleles confer cross-resistance to TDI-11055, which has previously been shown to act through an on-target mechanism of action by the discovery of drug-resistant *ENL* mutations.^10^ This is further supported by the known genetic and physical interactions between the PAF complex and the super elongation complex.^11,16,38–42^ Altogether, our findings emphasize the on-target nature of the cellular effects elicited by SR-0813, further validating it as a useful chemical probe to study ENL biology in acute leukemia.

## Data availability

All proteomics data have been deposited to the ProteomeXchange Consortium via PRIDE repository under dataset identifier PXD066867. Searched and averaged replicates of proteomics data can be found in supplementary information table: Table S6.

All 3’-UTR mRNA-sequencing data have been uploaded to GEO and can be found here GSE304765.

All code is available upon request.

## Acknowledgements

This work was supported by the Ono Pharma Foundation Breakthrough Science Initiative Awards Program, the Baxter Foundation Young Investigator Award, the National Institutes of Health (NIH) Office of the director (DP5-OD26380), and the National Cancer Institute (R01CA280720) (M.A.E). B.B.L is supported by the National Institute of Health (R35GM153476). The authors thank the Flow Cytometry and Genomics Core Facilities at The Scripps Research Institute for their guidance, access to flow cytometry instruments, and aid in sequencing experiments. We thank Dr. George Tsaprailis and his team in the Proteomics Core at the Herbert Wertheim UF Scripps Institute for their aid in running proteomics experiments.

---

**Supplementary Fig. 1.**
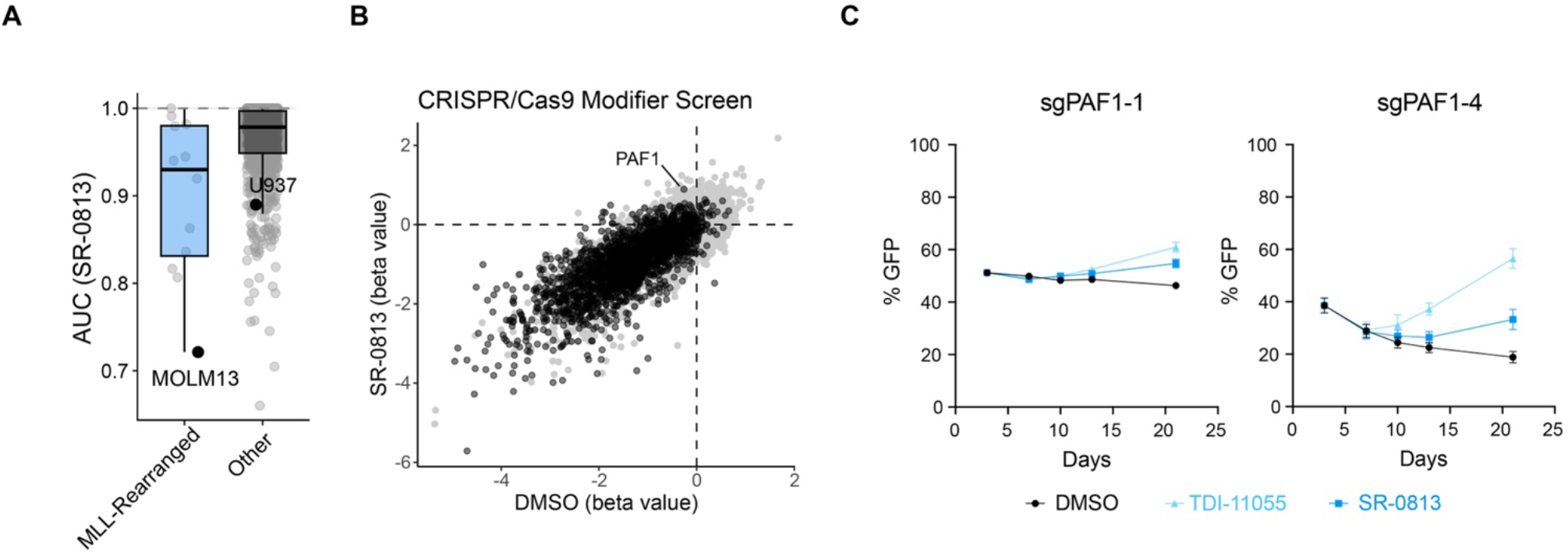
(A) PRISM Data of sensitivity to SR-0813 in cell lines harboring MLL-rearrangements. (B) Scatterplot of gene beta values following MAGecK MLE analysis of the two arms of the genome-wide CRISPR/Cas9 screen. Common essential genes (as determined by DepMap) are shown in black and others are shown in grey. (C) Competitive growth assay in MOLM13-Cas9 cells transduced with PAF1-targeting sgRNAs and treated with 10 µM TDI-11055, SR-0813, or DMSO over time.

**Supplementary Fig 2.**
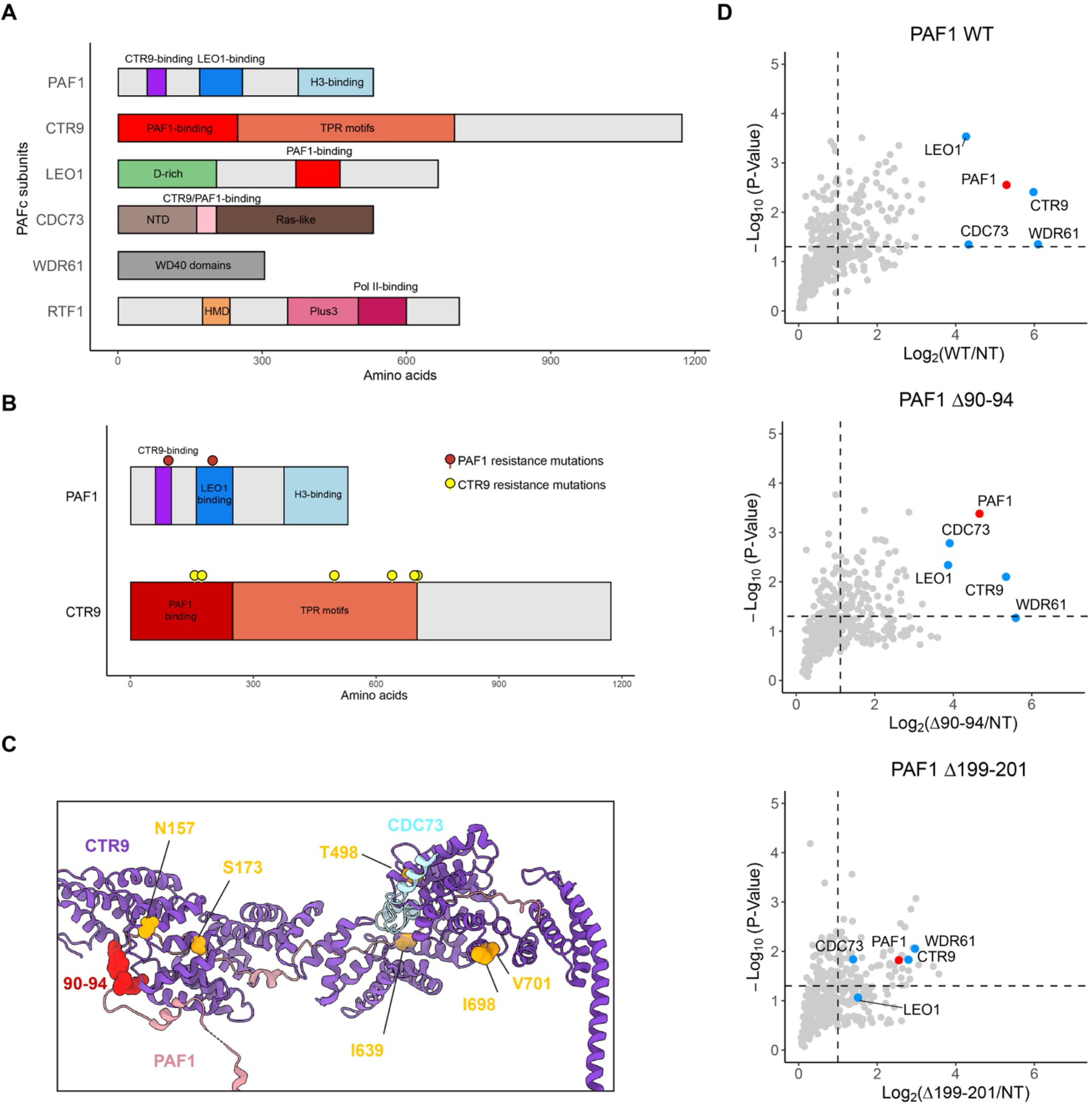
(A) PAFc protein domain maps adapted from Uniprot with PAF1 interaction sites highlighted. (B) PAF1 and CTR9 domain maps with resistance mutations highlighted in red and yellow, respectively (C) PAFc cryo-EM (PDB 6TED) depicting PAF1, CTR9 and CDC73 with amino acids 90-94 of PAF1 in red and sites of *CTR9* variants colored in yellow. (D) Volcano plots depicting PAF1 IP-MS relative to non-transduced controls. Data is filtered to remove protein values that are greater in the non-transduced (NT) control than in the pulldown, representing nonspecific binding.

**Supplementary Fig 3.**
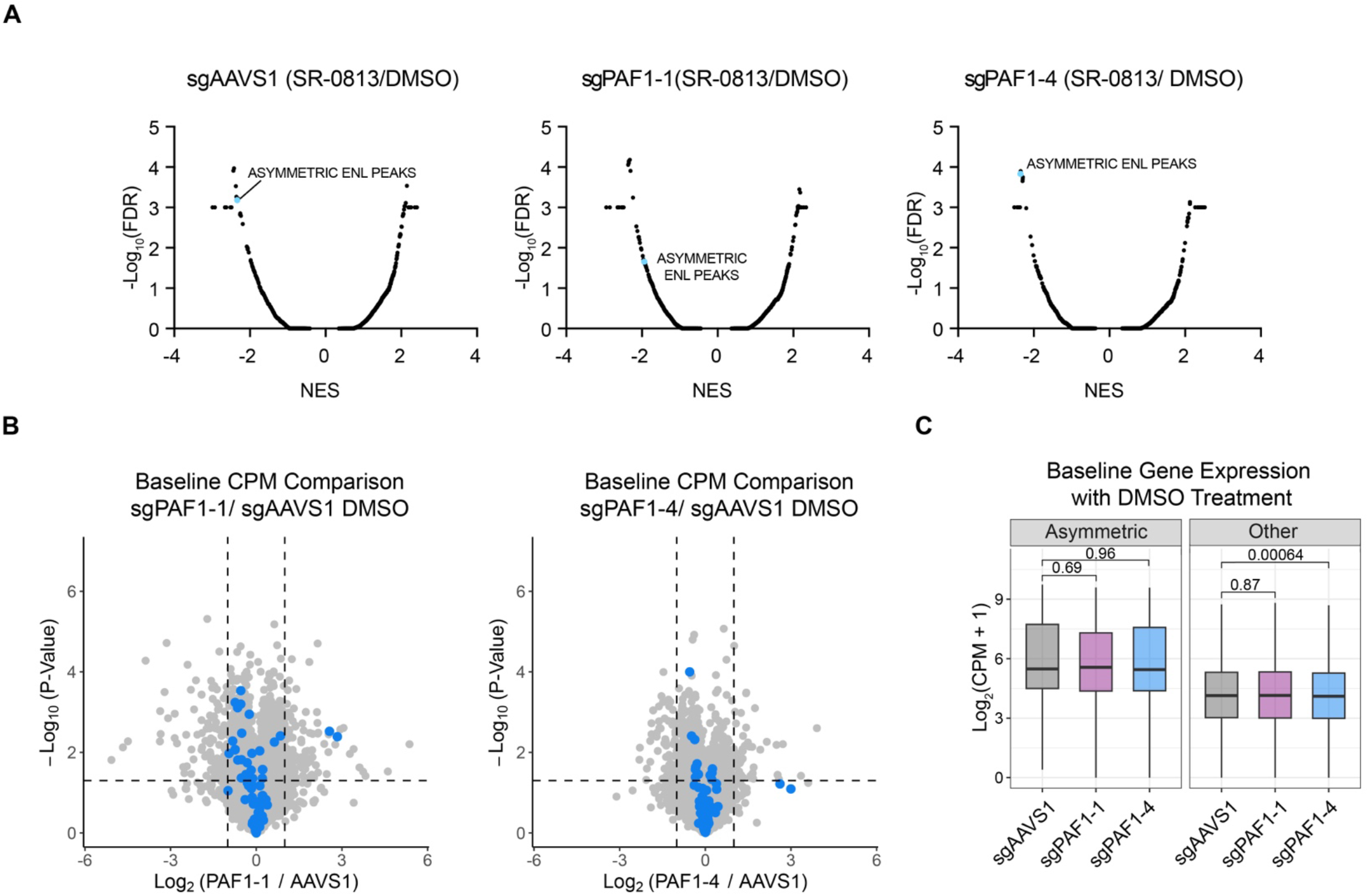
(A) GSEA analysis of gene expression changes following SR-0813 treatment using 3,297 gene sets in the Molecular Signatures Database (MSigDB) plus a custom gene set of asymmetric ENL targets. (B) Volcano plot comparing CPM transcript values for each cell line to the sgAAVS1 control group (DMSO-treated controls). Asymmetric ENL target genes are shown in blue. (C) Baseline CPM values from 3’-mRNA sequencing data in the DMSO-treated cell populations grouped by ENL target genes (asymmetric n=69, other n = 12702). *P*-values are calculated with a Welch’s two sample *t*-test.

